# Integrated Movement Models for Individual Tracking and Species Distribution Data

**DOI:** 10.1101/2024.06.19.599581

**Authors:** Frances E. Buderman, Ephraim M. Hanks, Viviana Ruiz-Gutierrez, Michael Shull, Robert K. Murphy, David A.W. Miller

## Abstract

1. While the quantity, quality, and variety of movement data has increased, methods that jointly allow for population- and species-level movement parameters to be estimated are still needed. We present a formal data integration approach to combine individual-level movement and population-level distribution data. We show how formal data integration can be used to improve precision of individual and population level movement parameters and allow additional population level metrics (e.g., connectivity) to be formally quantified.
2. We describe three components needed for an Integrated Movement Model (IMM): a model for individual movement, a model for among-individual heterogeneity, and a model to quantify changes in species distribution. We outline a general IMM framework and develop and apply a specific stochastic differential equation model to a case study of telemetry and species distribution data for golden eagles in western North American during spring migration.
3. We estimate eagle movements during spring migration from data collected between 2011 and 2019. Individual heterogeneity in migration behavior was modeled for two sub-populations, individuals that make significant northward migrations and those that remained in the southern Rocky Mountain region through the summer. As is the case with most tracking studies, the sample population of individual telemetered birds is not representative of the population, and underrepresents the proportion of long-distance migrants in. The IMM was able to provide a more biological accurate subpopulation structure by jointly estimating the structure using the species distribution data. In addition, the integrated approach a) improves accuracy of other estimated movement parameters, b) allows us to estimate the proportion of migratory and non-migratory birds in a given location and time, and c) estimate future spatio-temporal distributions of birds given a wintering location, which provide estimates of seasonal connectivity and migratory routes.
4. We demonstrate how IMMs can be successfully used to address the challenge of estimating accurate population level movement parameters. Our approach can be generalized to a broad range of available movement models and data types, allowing us to significantly improve our knowledge of migration ecology across taxonomic groups, and address population and continental level information needs for conservation and management.

## 1 Introduction

Our ability to address the effects of climate and land-use change on biodiversity and ecosystem health depends on our ability to predict how animal populations respond to changes in environmental conditions (Urban et al. 2016). To date, most forecasts of potential shifts in distributions are based on static patterns of habitat associations. However, an often ignored, yet critical, component for predicting these responses is the robust quantification of animal movement patterns (Nathan 2008; Flack et al. 2022).

The introduction of the global positioning system (GPS) in the early 1990’s began to revolutionize our ability to track wildlife. Tracking technology now allows us to collect large quantities of information about an individual with increasing precision and frequency for extended periods of time. This exponential increase in the availability of tracking data has coincided with an equally important advance in the statistical methods available for analyzing these data. These new methods have ranged in their focus, from estimating the true movement path given coarse data with measurement error (Buderman et al. 2015; Johnson et al. 2008a, 2008b; Jonsen, Flemming, and Myers 2005), to the size and location of the home-range of an individual (Fleming et al. 2015; Nilsen et al. 2008; Worton 1989) (e.g., MCP, KDE), and the preferential use of target resources on the landscape (Avgar et al. 2016; Manly, McDonald, and Thomas 2004; Thurfjell, Ciuti, and Boyce 2014) (RSF, SSF). Similarly, there is a growing interest in estimating underlying, or latent, behavioral states based on movement quantities related to locations of an individual through time, typically referred to in the movement literature as hidden Markov-models (McClintock et al. 2012). These advances in data collection and analysis have increased our understanding of the ecology and evolution of migratory behavior (Mueller et al. 2013; Gu et al. 2021), conservation of at risk species and populations (Liang et al. 2023), connectivity of migratory populations (Alheit and Bakun 2009; Kot et al. 2022), optimal habitats to prioritize for conservation (Yi et al. 2022), movement of pathogens (Takekawa et al. 2023), and ways in which species respond to climate change (Youngflesh et al. 2021; Horton et al. 2023).

Despite the amazing progress in the field of movement ecology, there still are significant challenges to continuing to advance our ability to better quantify movement. Movement behavior is a multi-scale, adaptive response that is influenced by both biotic interactions and abiotic environmental factors (Nathan et al., 2008). Quantifying movement is especially complex for migratory species, due to variation among multiple subpopulations within the species range, each of which makes unique movements that span continental and global scales. As a result, individual tracking data still have fundamental limitations when the goal is to understand movement behavior across the entire distributional range of a species (e.g., continental).

Individual tracking data are typically limited to relatively small sample sizes of tagged individuals and are almost always collected on a non-representative spatial subset of animals, relative to the entire range of the given species. Unfortunately, the cost and effort required to tag a balanced and representative sample of the variation in animal movement decisions is unattainable for all but a handful of range-restricted or non-migratory species. Thus, while tracking data provides critical fine-scale insights into movement behavior of individuals, they are not collected in ways that make inference about the movement and dynamics among multiple subpopulations feasible, simple, or straightforward. The consequence of these challenges is that continent- or region-wide insights regarding animal movement patterns are lacking for most at-risk species, and rarely are formally integrated into conservation planning. Overall, advances in this field are thus limited by our current inability to quantify and scale individual variation in movement behavior to population levels.

One potential solution to address the challenge of scaling from individuals to population level movements is to simultaneously quantify the global distribution of a species in space and time, as well as the variation in movement behavior across individuals and subpopulations. The relationship between individual behavior and population patterns has long fascinated ecologists (i.e., Turchin 1997; Wilson, Hanks, and Johnson 2018). For decades, mathematicians and biologists have considered scaling up individual movement models to population level models. Early work on this includes the work of Turchin (1998), with more recent work including that of Wilson et al., (2018) and Potts and Borger (2023). In general, these approaches do not jointly model both individual and species distribution data, which are often derived from occurrence data. Instead, they are either solely mathematical exercises (i.e., Turchin 1997) meant to motivate population level models, often taking the form of differential equations – or – studies of individual tracking data (i.e., Wilson et al., 2018) with the additional goal to scale results from a study of individual data up to an understanding of the resulting long-term patterns that would arise from species distributions. Hierarchical modeling approaches have also tried to link population-level processes and individual behavior (Scharf and Buderman 2020). However, inference is still limited to descriptions of the average behavior of the population of sampled individuals with transmitters. These methods do not jointly model independent sources of species distribution data with individual tracking data, and thus do not facilitate the combination of these two modern data streams.

One solution to fully scale individual-level data to the global, population-level scale is to integrate fine-scale tracking data with information about the global spatio-temporal distribution of the species and how it changes through time. Fine-scale estimates of intra-annual changes in species distributions have only recently become feasible due to the large-scale collection of species occurrence data and the development of temporally varying species distribution models. New platforms such as eBird (Sullivan et al. 2009), iNaturalist, and many others have empowered data collection with the help of thousands of volunteer contributors. Along with historic records and systematic surveys, these data form the backbone for efforts to quantify species distributions. Species’ distribution models are commonly used to assess species status, quantify patterns of diversity, and understand ecological drivers affecting species. However, aside from a few applications (e.g., Supp et al. 2015; Fuentes et al. 2023), records of species occurrences and associated species distribution models have not been seen as a source of movement data. These data can provide key information about the aggregation of individual movements throughout the year and have the potential to bridge gaps in individual tracking data that are collected non-representatively across the entire range and full annual cycle of a species. Formally integrating individual movement data and population-level, species distribution patterns will require new analytical methods that can link the two scales of inference throughout a species’ range and throughout the full annual cycle.

Our goal is to formalize a framework for the integrated modeling of species distributional data and individual tracking data to scale individual-level patterns to population-level processes across a species range and full annual cycle. Following the work of others, we define formal data integration as harnessing multiple data types to simultaneously estimate a common underlying state variable or process (Miller et al. 2019; Zipkin et al. 2019). The key here is that two types of data are used to estimate a common set of parameters, using a single integrated estimation framework. Other examples in the ecological sciences of formal data integration include models to estimate demographic processes, species distributions (Dorazio et al. 2012; Miller et al. 2019; Fletcher et al. 2019), population and community dynamics (Doser et al. 2022), and other ecosystem processes (Zipkin et al. 2021). These efforts have addressed how data are shared to make joint inference (Pacifici et al. 2017), methods to account for observation uncertainty and heterogeneity in survey effort (Dorazio et al. 2012; Miller et al. 2019; Zulian et al. 2021), and robust methods for cross-validation and designing optimal survey efforts for ecological models (Reich et al. 2018; Zulian et al. 2021).

To this point, attempts to combine inferences from individual tracking data and distribution data have not fully leveraged the statistical advances made in each field because they do not formally integrate the two data-sources (e.g., McCabe et al. 2021; Meehan et al, 2021). This leads to limitations such as failing to leverage information on speed and tortuosity available from tracking data or only using species distribution data to train or validate an individual-based movement or flow models (e.g., Fuentes et al., 2023; Tonelli et al., 2023). We describe the components of a formal integrated model for combining individual tracking and temporal distribution data. We then present an example of a hierarchical model that follows this general framework, and we illustrate its use through an integrated analysis of golden eagle (*Aquila chrysaetos*) tracking and species distribution data. Our approach addresses the current limitations in quantifying population-scale movement processes allowing us to facilitate spatiotemporal inference regarding how, when, and why animals move across large spatial and temporal scales.

## 2 Materials and Methods

### 2.1 General Framework for Integrated Movement Models

To fill this need, we propose a framework for integrated modeling of individual tracking and species distribution data. As we argue in the previous section, combining these data types to simultaneously estimate individual and population level movement patterns can unlock additional information and improve our inferences. Our ability to develop an integrated estimator requires three important statistical components:

1. **A movement model for individual tracking data.** This statistical model should allow for inference on processes of particular interest for the study and be appropriate for the data collected. Modern models for telemetry tracking data can capture autocorrelation in such data (Eisenhauer et al., 2022, Hooten et al., 2017, Johnson et al., 2008, Russell et al., 2018), varying levels of accuracy in the telemetry observations (Brost et al., 2015; Jonsen et al., 2020), and changing behavior over time (Eisenhauer et al., 2022; Glennie et al., 2022; Hanks et al., 2015), among other features. It is critical for integrated movement modeling that the statistical model chosen should be general enough to capture the range of behavior present in the whole population being modeled. This could be done either through a suite of related models, or a singular model with enough flexibility to model variation in movement behavior across the whole population.
2. **A model for the among-individual heterogeneity in movement behavior within a population.** Behavior of individuals within and across populations of the same species is often highly variable, some of which can be explained by measurable factors, such as age and sex, but a large proportion of which is not directly explainable by using easily measured features. In migratory species, two individuals with the same measurable characteristics may exhibit very different movement behavior over the full annual cycle; for example, one might migrate early while the other migrates late, or they may migrate to two different breeding grounds but share an area during winter. Our ability to quantify this variation in behavior is critical to improving our understanding of migratory connectivity (driven by variation in movement to and from breeding grounds), the landscape of risk (driven by variation in timing and routing of migrations), energetic tradeoffs (driven by migratory timing and habitat selection along the migratory route), and other behaviors. Without individual-level tracking data, it is in many cases impossible to identify multiple behavior patterns that result in the observed population-level pattern. Accounting for multiple movement behavioral patterns in individual tracking data is typically done by incorporating random effect modeling (i.e., Hooten et al., 2016; Scharf and Buderman 2020). However, an alternative and appealing framework that has rarely been used in movement modeling is a discrete mixture model, in which individuals can belong to different modes of movement behavior (i.e., Mastrantonio 2022, Eisenhauer et al. 2022).
3. **A movement model that quantifies changes in the species distribution**. An integrated model for individual and species distribution data requires a statistical model for the species distribution data that can predict changes in relative abundance at high spatial and temporal resolutions, and that is formally linked to the population level movement model arising from the individual model for telemetry data. In particular, the model for species distribution data should be a function of the same parameters that control the individual telemetry data model (from #1 above), or the scaled-up population model (from #2 above). This formal link – where parameters are shared and simultaneously estimated from information in both data types – is what defines an integrated movement model.

Together, these three models form an integrated model for both species’ distribution data and individual tracking data, with a shared process that captures various movement at both the individual level (with heterogeneity) and the population level. One must then be able to estimate shared parameters that govern movement and population dynamics. In many cases, models that are needed to capture heterogeneity in individual movement behavior across a population will be hierarchical in nature, and thus Bayesian approaches to estimation will often be the most straightforward for integrated movement modeling. We now propose one general class of movement models that can serve as the basis for integrated movement modeling for a relatively wide range of animal systems. We will develop a general modeling framework for integrated movement modeling, starting from a model for individual movement and moving up to the population level. This approach is illustrated in Figure 1. The specific models for telemetry, subpopulation structure, and species distribution data are kept general in this section, but we define them specifically for our case study in the next section.

**Figure 1:**
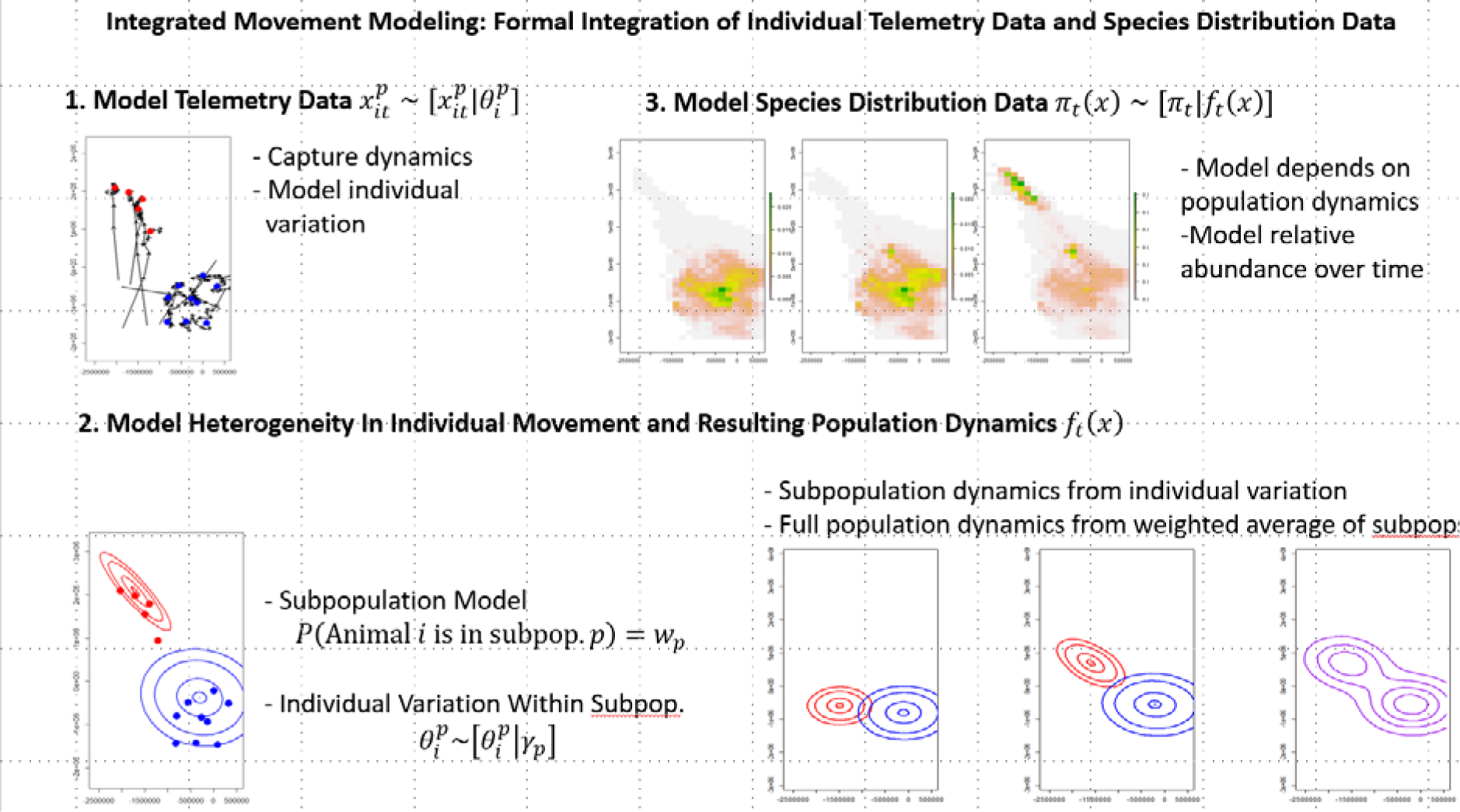
The Integrated Movement Modeling (IMM) Framework involves (1) modeling individual telemetry data, (2) specifying a model for how individual movement behavior varies within the population, and then deriving the population dynamics that result from these models, and (3) modeling species distribution data conditioned on these population dynamics.

To model heterogeneity among individuals in a population, we consider a mixture model approach in which there are multiple groups, or subpopulations, of animals. In principle, these subpopulations can differ in any aspect of movement behavior (i.e., subpopulations could be defined by the location they originate from), and individuals within a subpopulation can also have different behavior (i.e., onset of migration may differ among individuals).

Let 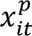 be the telemetry location for the *i*^th^ animal in subpopulation *p* at time *t*. We propose a general model for this data to be

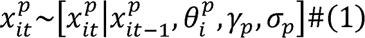

As is standard in movement modeling, we model animal locations conditional on previous locations in time (here we consider a Markovian model, conditioned only on the previous observed time) and also dependent on individual specific parameters 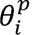, subpopulation parameters *γ_p_*, and variance (observation error) parameters σ*_p_*. This model form is general enough to encompass integrated step selection function models (e.g., Avgar et al. 2016), Markov chain models (e.g., Wilson et al. 2018), many standard stochastic differential equation models (e.g., Preisler et al. 2004; Preisler et a. 2014; Eisenhauer et al. 2022), and others.

To model variation among individuals, we consider a general model in which the individual-specific parameters depend on the subpopulation-level parameters

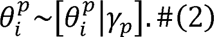

For example, this could be a standard random effects model in which the individual-specific movement parameters are normally distributed around a shared subpopulation mean. Together with a model for how individuals are structured into these subpopulations,

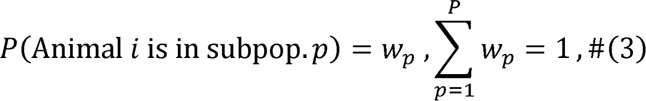

we have a complete model for a population of animals with individual and subpopulation heterogeneity in movement behavior.

We now consider a model for the population dynamics of the spatio-temporal distribution of this population, built probabilistically from the individual model and subpopulation structure above. Assume that at time *t* = 0 that each subpopulation’s spatial distribution is defined by a general spatial distribution

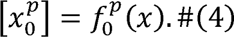

Then under the assumption that this initial distribution is independent of (2) above, the population dynamics can be obtained by marginalizing over individual and subpopulation dynamics in the individual movement model (1)

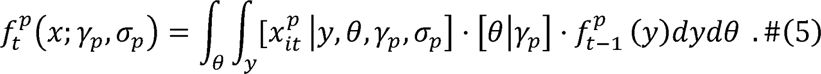

In our example analysis, we show one approach that allows this integration to be done analytically. This is a discrete time model for the dynamics of a subpopulation of animals moving under the movement model, together with individual heterogeneity, proposed above. The result is 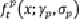, the spatial distribution of the *p*^th^ subpopulation at time *t*, conditioned on an initial spatial distribution and the population level parameters governing individual movement. Then the full population distribution is just the weighted sum of these subpopulation distributions

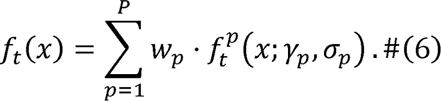

This full spatio-temporal population distribution can then be used as the mean in a model of the observed species distribution data

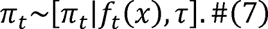

Where τ, are parameters related to the observation error (i.e., variance and/or correlation) of the observed species distribution data {τ_1_,τ_2_,…,τ_T_}. Together these models provide a formal approach for integrated individual and species distribution data.

### 2.2 Case Study: Golden Eagle Spring Migration in the Western North America

We now illustrate this framework for formal integration of individual tracking and species distribution data through an analysis of spring migratory behavior of golden eagles that had been tagged with satellite transmitters while in the conterminous western United States (U.S.).

The golden eagle has a Holarctic distribution, occurring throughout most of North America, Europe, Asia, and parts of northern Africa (Katzner et al. 2022). In the eastern U.S., golden eagles have been extirpated as a breeding species but individuals from eastern Canada regularly overwinter (Katzner et al. 2020). Currently, numbers of golden eagles in the western U.S. tentatively are stable but likely will decline as anthropogenic mortality increases (Millsap et al. 2022). Golden eagles in the U.S. are protected under the Bald and Golden Eagle Protection Act (BGEPA; 16 U.S.C. 668-668c), which prohibits take, defined as “pursue, shoot, shoot at, poison, wound, kill, capture, trap, collect, molest or disturb,” unless the take is incidental to otherwise lawful activities authorized by permit from the U.S. Fish and Wildlife Service (USFWS). Examples of situations where incidental take may occur but possibly be authorized by permit are collisions with turbines at wind energy projects (Beston et al. 2016) and electrocutions on power line poles (Mojica et al. 2018). Take of a wild golden eagle must by offset by saving or creating (e.g., by improving reproductive success) 1.2 golden eagles elsewhere; the mitigation offset ratio is greater than equivalent due to the golden eagle’s tentative population status. However, the number of golden eagles that can be taken under permit is strictly limited, at two spatial scales: the Eagle Management Unit (EMU; large scale, each unit encompassing about one-third of coterminous U.S. states) and Local Area Population (LAP; area within 174 km of the site or activity to which a permit applies) (U.S. Fish and Wildlife Service 2016). Dynamic movement of golden eagles across these spatial scales is not well understood and could influence estimates of the eagle’s population size within these areas. Utility of the coarse-scale EMUs for determining upper thresholds for permitted take is uncertain; interconnected subregional populations may exist that could better serve as single population management units for incidental-take permitting decisions.

To inform golden eagle management decisions including limits on permitted take, the USFWS worked with a network of collaborators to amass telemetry data from >600 golden eagles tracked via satellite or GSM (Global System for Mobile Communications) transmitters in the conterminous western U.S. (Millsap et al. 2022). Main objectives of the work were to estimate annual survival and identify major causes of mortality. Subsets of the data have been used to address other management information gaps, such as movement behavior, e.g., identifying major migration corridors (Bedrosian et al. 2018, Brown et al. 2017), describing non-routine, long-distance movements (Poessel et al. 2016), and documenting juvenile dispersal (Murphy et al. 2017) and natal dispersal distance (Murphy et al. 2019). However, these individual-level results are difficult to scale to the entire population, or to sub-populations such as EMUs and LAPs, and do not provide detailed information on the relative number of individuals that are being exposed to risk, or the relative importance of identified migratory pathways, stopover areas, or wintering grounds. To fill these information gaps, the Cornell Lab has closely collaborated with the USFWS Division of Migratory Bird Management’s National Raptor Program to validate and integrate eBird relative abundance information for golden eagles, to inform population size estimates and policy germane to incidental take permitting. No reliable framework exists for integrating these sources of information to ascertain relative exposure risk along migratory pathways and stopover areas, or to identify which subregional populations are interconnected such that they could be managed as single population management units for incidental-take permitting decisions.

#### 2.2.1 Golden Eagle Data Description

We aggregated telemetry data from 136 golden eagles tracked by the USFWS via satellite telemetry between 2011 and 2019; individual eagles in our sample were represented by data from 1-8 spring migration seasons each. Eagles were tagged in the Colorado Plateau, Rocky Mountain (south of Montana), Central Great Plains, Southern Great Plains, and Texas Trans-Pecos regions, encompassing roughly the eastern one-half of the species’ range in the coterminous western U.S. Most (77.9%) were tagged with satellite transmitters when they were large (7- to 8-week old) nestlings; these permanently dispersed from natal areas by the end of their first year of life (Murphy et al. 2017). Others (22.1%) were trapped and tagged when in their second year of life or older; including some that were settled on breeding territories when ≥4 years of age (Murphy et al. 2019). Transmitters were solar Argos/GPS 45-g and 70-g platform terminal transmitter units (Microwave Telemetry, Inc., Columbia, MD); each was attached in a backpack configuration via “Y-harness” constructed of Teflon ribbon (Murphy et al. 2017). Transmitters collected GPS locations hourly each day during at least 0900-1600 H; PTT location accuracy was ± 19 m. Our dataset for a given eagle included a single “daily” location for each 24-hour period, derived by averaging all GPS locations available for the period. In this analysis, we consider only the spring migration, and thus subset each available year of telemetry data to the time between Julian day 50 (February 19) and Julian day 105 (April 15). While this window does not capture the entire migratory season for the species, it does contain most of the migration detectable in the telemetry data as well as the eBird species distribution data.

As we are considering just the spring migration, we treated each year of data from a bird as being independent of all other years of data for that individual (hereafter, each sample will be referred to as a bird year). We also defined each bird year as being part of one of two subpopulations, with the first subpopulation being all bird-years where the bird did not pass north of the 51-degree Latitude line and the second subpopulation being all bird years where the bird moved north of this latitude within our spring window. This resulted in 336 bird years in the first subpopulation and 18 bird years in the second subpopulation. These daily telemetry data are the individual telemetry data we will use in our Integrated Movement Model. Figure 2 shows daily telemetry data, with arrows pointing to successive days for each bird. Figure 2a shows all tracked birds in the first subpopulation (those that did not cross the 51-degree latitude line) and Figure 2b shows all tracked animals in the second subpopulation (those who did cross north of the 51-degree latitude line).

**Figure 2:**
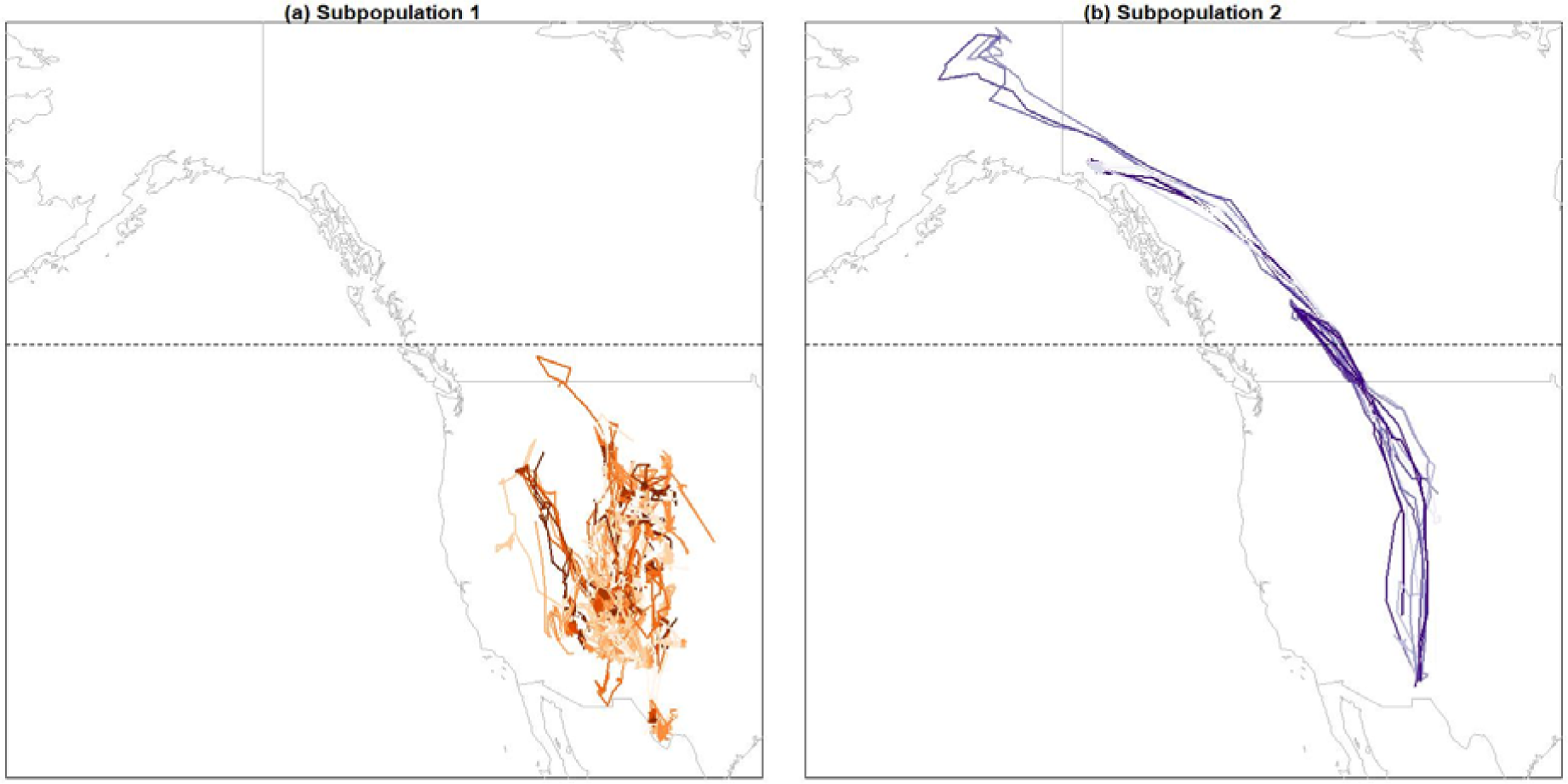
Daily spring GPS locations for 136 golden eagles in the western U.S and Canada, with lines connecting sequential locations. The birds are divided into two subpopulations: (a) those that spend the spring (Julian day 50 (February 19) and Julian day 105 (April 15) south of the 51-degree latitude line (dashed line) and (b) those that move north of the line. Note that colors repeat due to the number of individuals.

For species distribution data, we used eBird relative abundance information from 2019 (Fink et al. 2019, 2021). The Cornell Lab of Ornithology has developed the Adaptive Spatio-Temporal Exploratory Model (AdaSTEM; Fink et al. 2014; Fink, Damoulas, & Dave 2013), which processes huge numbers of individual citizen-science records, accounts for spatial heterogeneity in sampling effort, observer skill, and rarity of species to estimate the relative abundance of a species over space and time (Fink, Damoulas, & Dave 2013). This eBird relative abundance information is available at a 2.8×2.8km resolution, for each week of the year, throughout the entire distributional range of a given species. We subset the relative abundance down to just the western U.S., aggregated to 100-km resolution, and normalized the gridded relative abundance to sum to unity. This weekly, normalized, eBird relative abundance is the species distribution data we will use in our Integrated Movement Model. Figure 3 presents this weekly species distribution data, showing the partially-migrating nature of golden eagles, with some of the population migrating north to Alaska and northwestern Canada, while some of the population remains in the western U.S.

**Figure 3:**
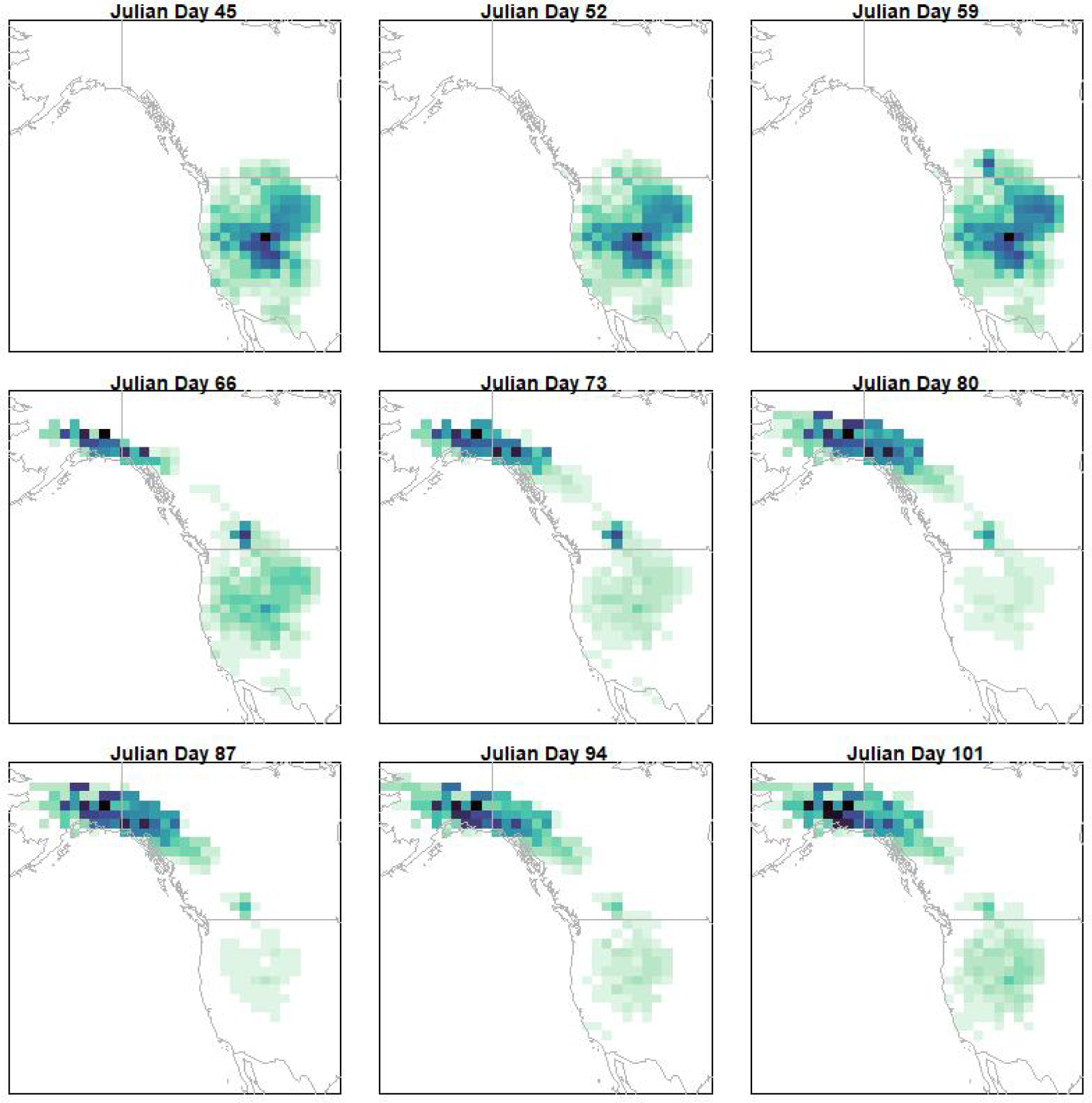
Weekly distribution of golden eagles in the western U.S. and Canada during spring 2019, obtained from eBird status and trends.

#### 2.2.2 A Hierarchical Integrated Movement Model for Golden Eagles

We now illustrate our integrated movement modeling framework from Eq. 1-7 on the spring telemetry and eBird relative abundance data of golden eagles. The class of individual models we will consider are stochastic differential equation (SDE) models, with movement governed by a potential function (Preisler et al., 2004; Eisenhauer et al., 2022; Russell et al., 2018). This class of models is particularly appealing for modeling migratory behavior because potential functions provide a straightforward approach to modeling movement along gradients or toward summer/winter ranges, however, as mentioned, other movement models could be used.

To model heterogeneity among individuals in a population, we consider a mixture model approach in which there are multiple groups, or subpopulations, of animals. In theory, these subpopulations can differ in any aspect of movement behavior, such as timing of migration, distance, duration, or number of geographic centroids; however, for this example we will focus on the location of geographic centroids. We will, for simplicity, assume that movement behavior is fixed over time; this does not mean that an individual’s locations are fixed through time, but that the parameters describing the subpopulation’s behavior do not change over time. In our example analysis in the following section, we consider only movement during the spring migration season – a time frame in which this assumption is appropriate.

##### 2.2.2.1 An SDE Model for Tracking Data

We will first define an SDE for individual telemetry data, which will correspond to Eqs. 1-3 in our general framework. Let 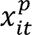 be the telemetry location for the *i*-th animal in subpopulation *p* at time *t*. We modeled movement using an SDE, with movement in a quadratic potential function centered around an attractive point 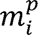 which is specific to animal *i.* Let

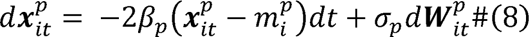

where 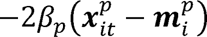 is the negative gradient of a quadratic potential function 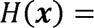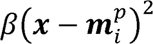 evaluated at the location of animal *i*. In this SDE, an animal’s mean movement at any given time is directly towards the attractive central location 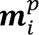, with random variation around that mean modeled using two-dimensional standard Brownian motion (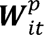)).

To make SDEs numerically tractable for statistical inference, we consider a discrete Euler-Maruyama approximation to the continuous time SDE (Eisenhauer et al., 2022; Russell et al., 2018). This results in the following time-discretized model, where Δ is the time-step between observations.

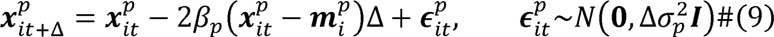

Variations of the above general framework have been used in multiple studies of animal movement to study a wide range of movement behaviors (Preisler et al., 2013; Eisenhauer et al., 2022; Russell et al., 2018).

This model is a model for individual telemetry data and corresponds to Eq. 1 in our general framework. For each animal there is an unknown attractive point 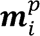, and for each subpopulation there are population level parameters *β_p_* and *σ_p_*. If we were fitting this model alone to the telemetry data, the unknown parameters could be estimated using ordinary least squares or maximum likelihood methods (Preisler et al., 2004; Russell et al., 2018; Eisenhauer et al., 2022). However, once we pair this model with a population-level model for the species distribution data, the resulting integrated movement model will be hierarchical and will be best fit using Bayesian methods.

Following Eq. 2 in our general framework, we can assume that the individual-level attractive points 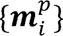 for individuals belonging to each subpopulation are located in relatively similar spatial locations They can then be modeled in a typical hierarchical fashion where the individual-level attractive points arise from a 2-dimensional Gaussian distribution with a shared mean and covariance

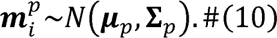

Allowing for multiple subpopulations is achieved by using a mixture of Gaussian distributions to describe the population; mixture models are a flexible framework and are not limited to just two mixtures of the same distribution. The mixture model would impose a statistical structure on Eq. 3 in our general framework.

##### 2.2.2.2 An SDE Model for Population Dynamics

We now develop a model for the movement (or diffusion) of a population of birds under the specific movement model in Eqs. 9-10. This corresponds to Eq. 4-6 in our general framework. If we assume that the initial distribution at time *t* = 0 of the *p-*th subpopulation is 2-dimensional Gaussian with

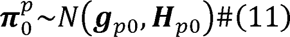

and additionally marginalize over the distribution of attractive points (10), then the distribution of the *p*-th subpopulation at time *t* can be found sequentially using standard multivariate normal distribution theory combining Eqs. (9)-(11)

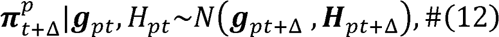

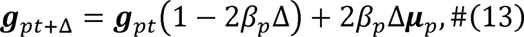

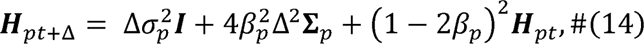

Under this model, at each successive time step, the subpopulation distribution moves from its initial distribution closer to a stationary distribution defined by the distribution of subpopulation attractive points (Eq. 10).

At any given time, we assume that the full population is a weighted sum of the subpopulation distributions (Eq. 7), with weights {*w_p_*} summing to 1

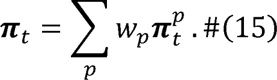

The population dynamics under this model assume a population of animals, divided into subpopulations, with individuals in each subpopulation having relatively similar spatial locations of attractive points (Eq. 10) and each individual moving in a quadratic potential function centered on their individual attractive point (Eq. 9). The population dynamics are extremely straightforward to calculate in this model, as at any given time the population distribution is a weighted mixture of 2-dimensional Gaussian densities with means {***g****_pt_*} and covariance matrices {***H****_pt_*} that can be deterministically calculated using Eqs. 12-14.

The key need for integrated movement modeling is a clear approach for scaling from individual to population level dynamics. In this example we address this need through a hierarchical random effects approach, which makes population dynamics very computationally efficient to compute and is one strong advantage of our probability-based approach for population dynamics (Eqs. 4-6). There are other possible approaches, such as scaling from individual SDEs to their population distribution Fokker-Plank equations (Gardiner 2009) or scaling grid-based movement models like step selection functions to population-level differential equations (Fricks and Hanks 2018; Potts and Borger 2022), but the approach we propose here is notable for the simplicity of the calculations required for the population dynamics.

##### 2.2.2.3 An SDE Model for Species Distribution Data

We now propose a model for species distribution data at a given time, which has a mean equal to the population distribution in (15). At time *t,* we assume that the species distribution data is in gridded form, with *sd_t_*(*x*) being the proportion of the population at the grid cell with center *x* at time *t.* We assume that the species distribution data is normalized, and thus sums to 1 over all spatial grid cells (i.e., relative abundance). We want a probability model for ***sd_t_*** = {*sd_t_*(*x*_1_),*sd_t_*(*x*_2_),…*sd_t_*(*x_M_*)}, the vector of observed species distribution relative abundances, with mean equal to the population distribution in (15). We propose a multinomial distribution for a scaled version of the species distribution data. Let *N* be a positive integer and let 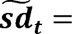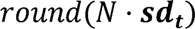 be a vector of closest integers to the scaled species distribution data. We propose modeling this quantity as

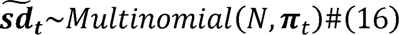

where π*_t_* is given by (15). In this model, the scaling factor *N* is best thought of as a dispersion or variance parameter, with smaller *N* leading to higher variance in the species distribution probabilities. This provides a model for species distribution data that has a mean given by the population model resulting from a population of individuals moving under our SDE model, with a flexible variance, *N*, that can capture an appropriate level of mismatch between the species distribution and our model. Together, (9)-(16) provide an example of a set of models that satisfy the requirements for an integrated movement model.

##### 2.2.2.4 Fitting the Integrated Movement Model

As the JMM described above is a hierarchical model, we chose to take a Bayesian approach to inference. For each subpopulation, we assign vague Gaussian and Inverse Wishart priors to the subpopulation-level mean and covariance of these movement centers:

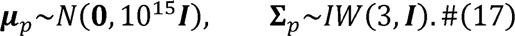

As the movement parameter *β* must be non-negative, we assign it a vague truncated normal prior, constrained to be greater than zero and assign an inverse gamma prior for the variance parameter σ^2^:

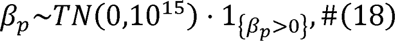

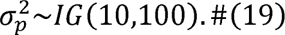

We assign a Poisson prior to the scaling (or dispersion) parameter *N* from the species distribution model, and vague priors to the initial subpopulation distribution means and covariances:

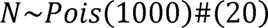

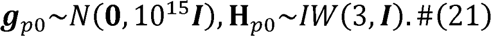

For the population weights {*w*_1_, *w*_2_}, we assign a uniform prior to *w*_1_, with the constraint that each population contain at least 10% of the population mass:

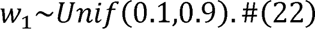

We fit this model using a custom Markov chain Monte Carlo sampler, coded in R, with adaptive tuning using the log-adaptive approach of Shaby and Wells (2010). We ran the sampler for 4,500,000 iterations and assessed convergence visually. This chain took under 48 hours to run on a single computer with a 16-core Intel I9 processor with clock speed 3.2GHz. Details on the MCMC algorithm have been provided in Appendix A. We also conducted a simulation study, by simulating both telemetry and species distribution data from the IMM model with all parameters set at the posterior means; details of this simulation study are in Appendix B and code and data to replicate the simulation are provided as supplemental material.

## 3 Results

Figure 4 shows results from fitting the integrated movement model to golden eagles in the western U.S. and Canada. Figure 4a-b shows posterior distributions for parameters β*p*, the subpopulation-specific strength of attraction to an individual-movement center, and 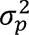, the subpopulation-specific variance parameter that captures both observation error and variation in observed locations around the mean movement in equation (9). These estimates reveal that birds in the north-moving subpopulation (*p*=2) are more highly attracted to their movement centers than are birds from subpopulation 1 but also show more stochastic variability in their movement. Figures 4c-d show the estimated distributions of movement centers (i.e., the estimated Gaussian distribution of 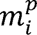 from Eq. 10, defined by the posterior mean estimates of the mean parameters µ*_p_* and variance-covariance matrix Σ*_p_*) for subpopulations 1 and 2, with subpopulation 2 showing movement centers distributed along a migration pathway to the Northwest of the initial species distribution mass. Figure 4e shows the posterior distribution of *w*_1_, the proportion of individuals estimated to be in subpopulation 1, and Figure 4f shows the posterior distribution for the scaling/dispersion parameter *N* for the species distribution data (see Eq. 16).

**Figure 4:**
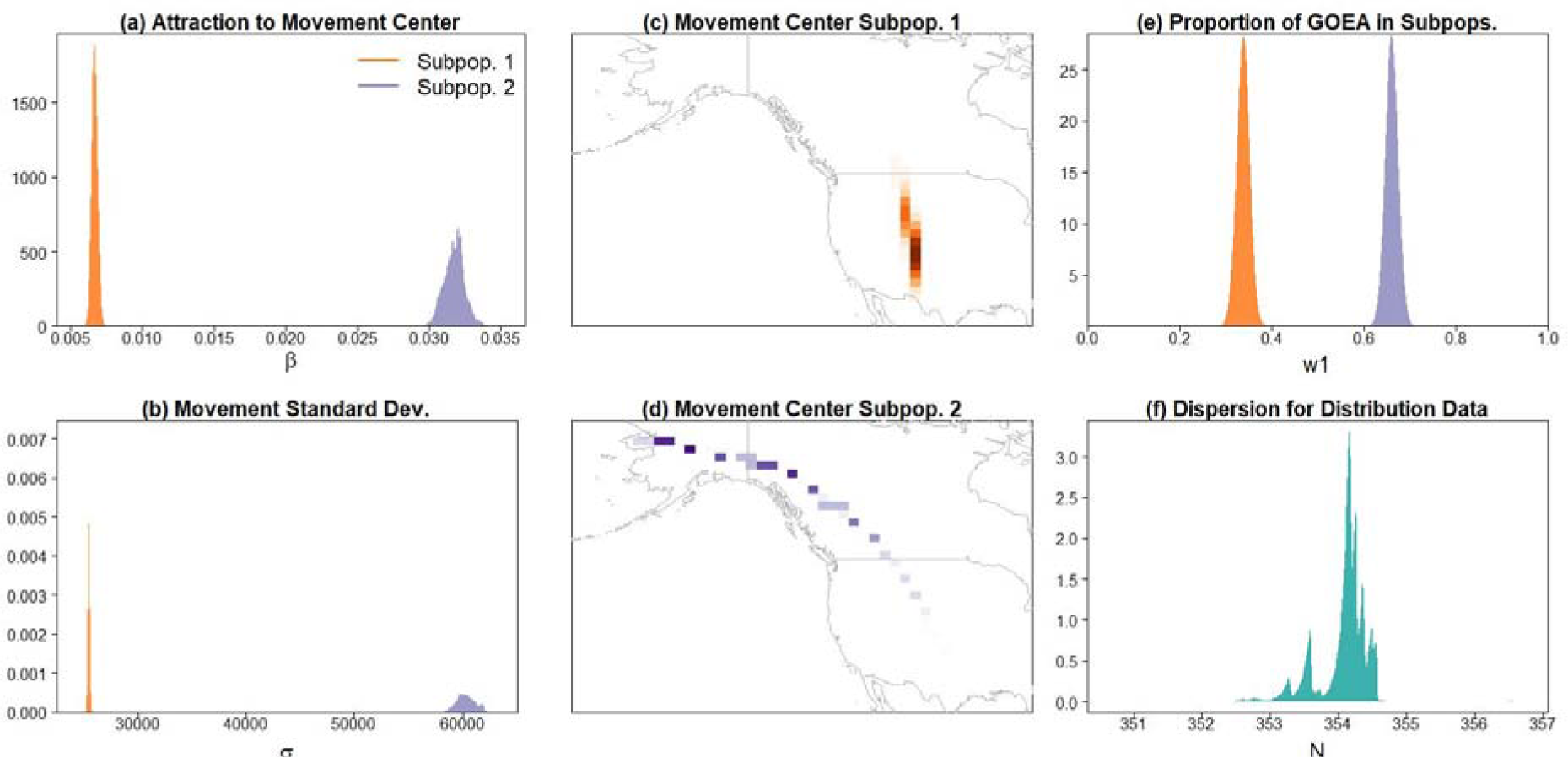
Posterior distributions from selected parameters from the integrated movement model analysis of the golden eagle datasets. Subpopulation-specific posterior means (a) and standard deviations (b) for attraction to individual movement centers. The movement centers 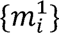 for subpopulation 1 are centered in the conterminous U.S. (c), while the corresponding movement centers for subpopulation 2 are spread along the migratory route to Alaska (d). We estimate 34% of the total population is in subpopulation 1 (e). The dispersion parameter *N* has posterior mean of 354 (f).

To illustrate the estimated subpopulation behavior, Figure 5a shows a triweekly subset of the eBird species distribution data and Figures 5b-c show posterior mean subpopulation distributions for both subpopulations (Eq. 12-14). Figures 5d-e show the pointwise posterior predictive mean percent of the observed eBird species distribution attributed to each subpopulation over time. This was calculated pointwise for each pixel, with the shown posterior predictive mean percent for the *g*-th pixel calculated as 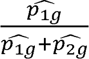, where 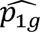 is the posterior mean subpopulation distribution for subpopulation 1, and 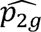 is the posterior mean subpopulation distribution for subpopulation 2. Figures 5d-e illustrate how the integrated movement model provides a formal approach for splitting an observed species distribution into multiple subpopulation distributions, each with their own dynamics informed by observed telemetry data.

**Figure 5:**
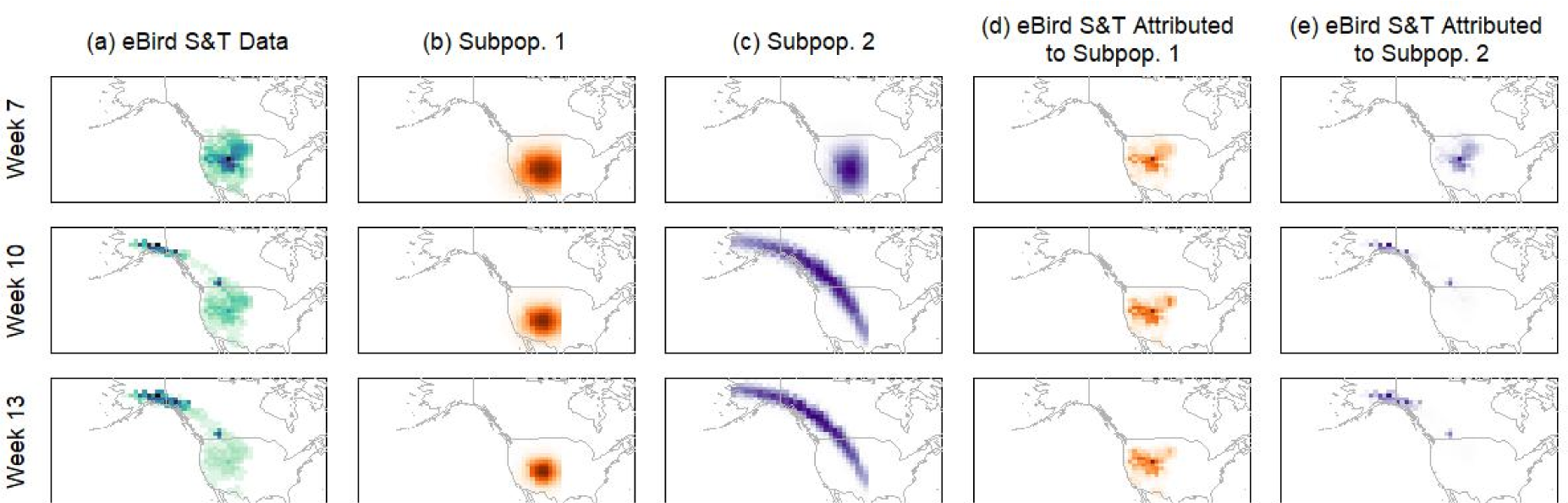
Observed biweekly distribution of golden eagles in the western U.S. during spring 2019 according to the eBird Status & Trends data (a1-4). Posterior mean of the distributions of subpopulation 1 (b1-4) and 2 (c1-4). Pointwise posterior predictive mean of the observed golden eagle distribution belonging to subpopulation 1 (d1-4) and 2 (e1-4).

In Figure 6 we illustrate how inference from the integrated model differs from a single data source alone. First, telemetry data are almost seldom collected in a way that is representative of the full population. In Figure 6a, we calculate the proportion of birds in our non-migratory subpopulation using just the telemetry data. This inference is done using a Bayesian approach with subpopulation membership modeled as a Bernoulli random variable with proportion *w*_1_ of birds in subpopulation 1 having the same prior (22) as in our integrated analysis. As the telemetry data have a large proportion of non-migrating birds, the estimated proportion of birds in this subpopulation is over 95%. However, once we jointly model the telemetry data with the species distribution data, the estimated proportion in subpopulation 1 is 34% in our study region. Similarly, in Figure 6(b) we compare estimates of the subpopulation movement parameters β_1_ and β_2_ from the telemetry data alone, obtained by fitting the model defined by Eqs. 9-10 and 17-18 with comparable estimates from the full integrated model. The inclusion of species distribution data results in estimates of lower attraction to the individual movement centers – this is especially evident in subpopulation 1.

**Figure 6:**
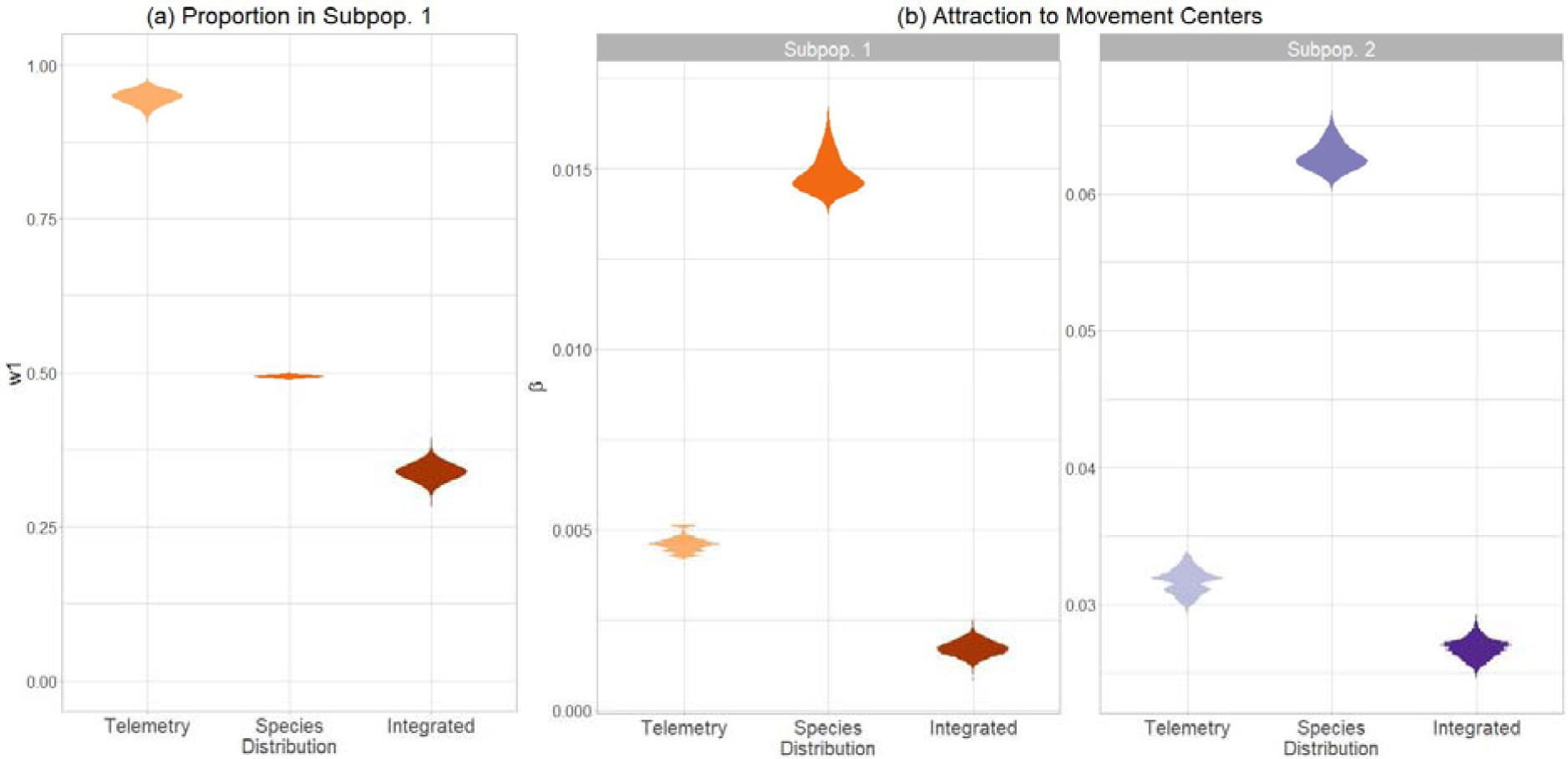
Comparison on the percentage of birds assigned to the migratory subpopulation is misrepresented by the typical biased sampling of animals in telemetry studies (a) and the attraction to individual movement centers for each subpopulation (b) using just telemetry data, just species distribution data, and the integrated movement model.

Figure 7 shows full probabilistic predictions of where animals who winter in a defined spatial location (i.e., a management unit) will occur over the course of the spring migratory season, something that would be impossible without an understanding of the spatial distribution of subpopulations such as we obtained from our integrated analysis. Figure 7 shows the posterior mean spatio-temporal distribution of the location of an animal beginning in central New Mexico (shown by the black “X”) over the temporal range of our study. The predictions result from using the estimated spatial proportion of birds in each of the two modeled subpopulations at the marked location (see Figure 5e1) to weight the dynamics of each subpopulation, given that birds in that subpopulation start in the marked location. This is accomplished through application of the dynamics in equations (12)-(14), using posterior mean values for all parameters except for the initial distribution parameters {*g*_10_, *g*_20_, *H*_10_, *H*_20_) which are chosen so that more than 99% of the entire initial subpopulation resides within 100km of the starting location marked by the black “X” (by fixing *g_p_*_0_ to be the center of the grid cell and *H_p_*_0_ to be a diagonal matrix with diagonal entries chosen so that the edges of the grid cell correspond to the 0.005 and 0.995 quantile of the marginal Gaussian distribution in both the latitude and longitude directions). The predictions show that a large number of birds are predicted to remain near the initial location, with a smaller, but still significant, number of birds migrating North. In Figure 8 we show posterior mean estimates of the proportion of birds in a given spatial location (shown by purple points) that are in subpopulation 2, the more migratory subpopulation. In the light point in Canada, we estimate a large early pulse of migrators moving through the location, while the dark point in Montana shows an increasing proportion of migrators through the spring migration.

**Figure 7:**
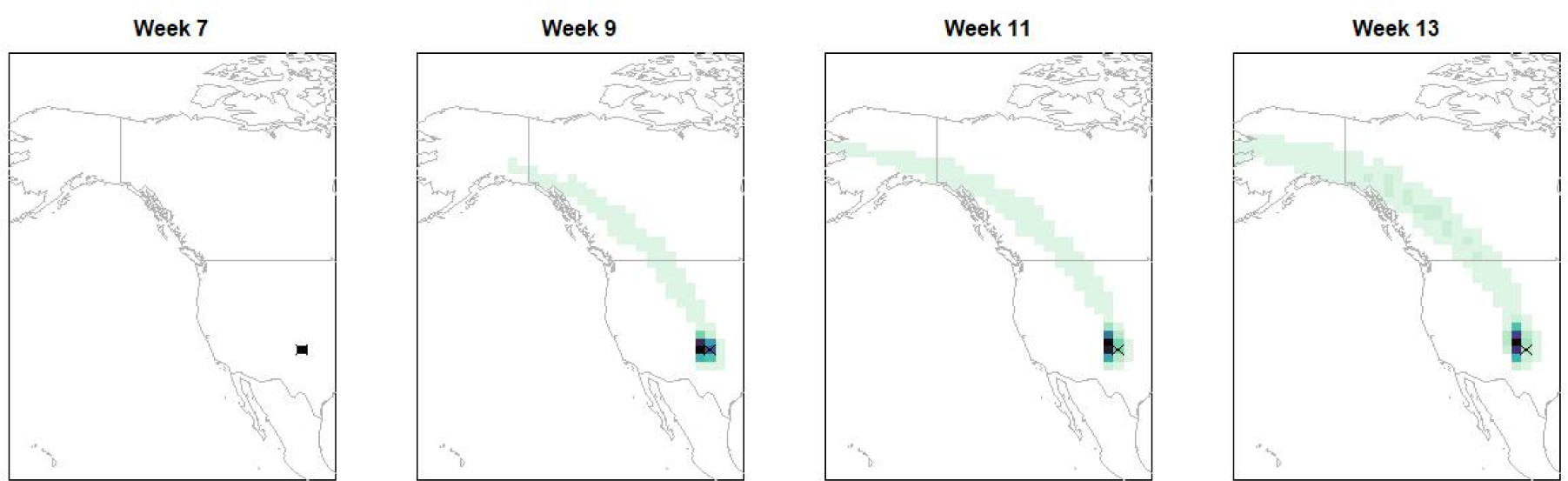
Predictions of the spatial distribution of the population of birds beginning at the grid cell marked with an “x” through spring migration.

**Figure 8:**
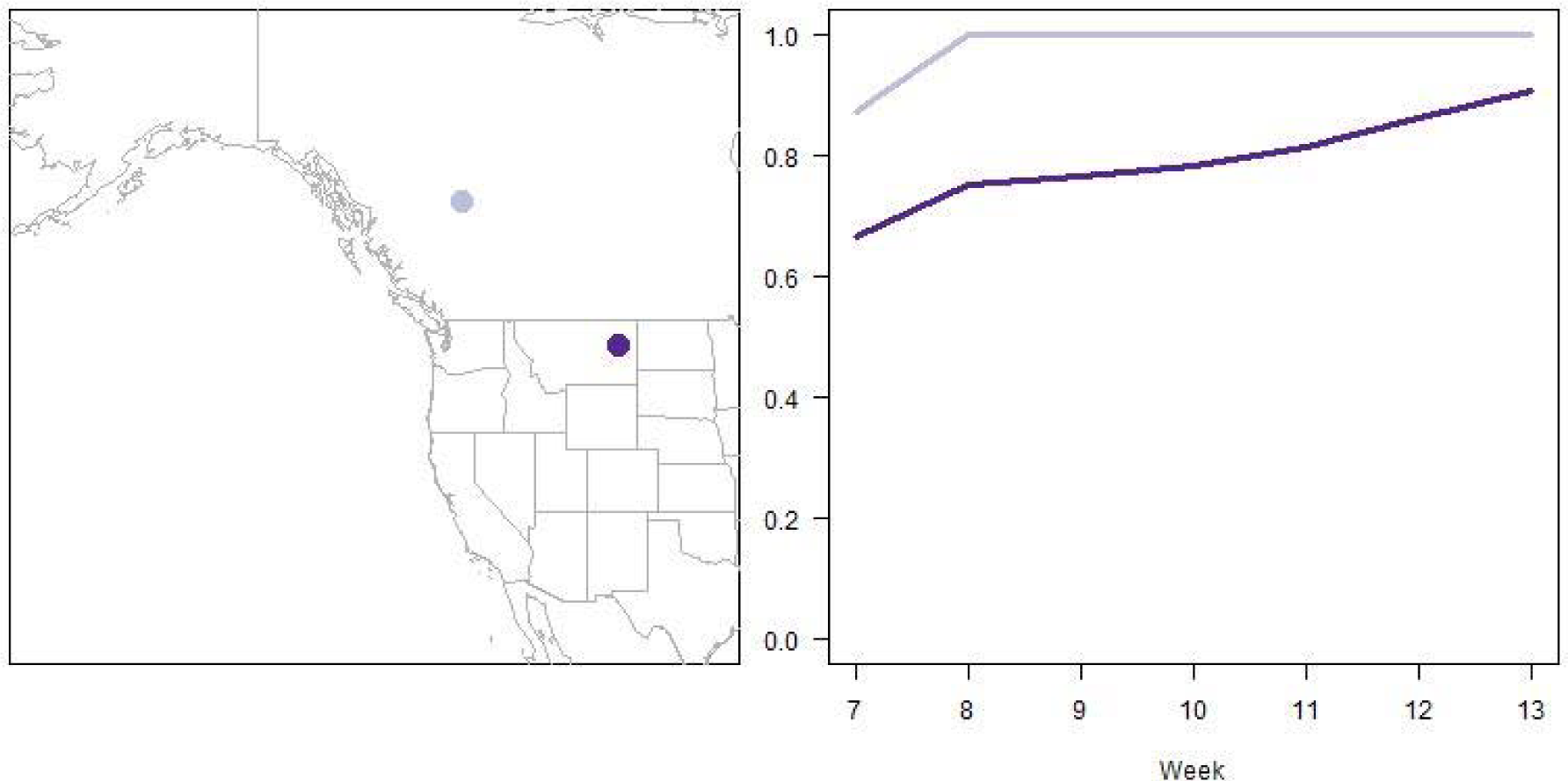
Proportion of individuals in subpopulation 2 though spring migration at two different spatial locations (b).

## 4 Discussion

In this manuscript, we demonstrate a general framework for Integrated Movement Modeling that combines both individual telemetry and species distribution data. We show how this approach addresses the challenge of simultaneously quantifying movement dynamics at the level of whole species’ distributions while accounting for variation in movement behavior across individuals and subpopulations. By applying this approach to spring migration of golden eagles in the western U.S, we demonstrate that the joint inference from both individual and species distribution data has benefits that are not possible when making inference from just one data stream.

There are several main takeaways from our modeling approach that come from the golden eagle case study. First are the unique results that are obtained from the integrated movement model, which contrast those available from each dataset individually. Foremost is the ability to directly estimate population-level parameters including the overall proportion of migratory versus non-migratory individuals within the modeled range. For golden eagles, we estimated that about 34% of the golden eagle western population belongs in subpopulation 1, which is generally stationary throughout the migratory season. However, we would have estimated that about 95% of the population is in subpopulation 1 if we had based our inferences solely on telemetry data, a sample we know is not randomly chosen from the whole population. This occurs because a disproportionate number of telemetered individuals remained south of 51-degrees latitude, which is inconsistent with seasonal changes in the distributional range of the species (Figure 3).

We are also able to uniquely generate fine-scale estimates of vital population level processes, such as the proportion of individuals in a given location that belong to a subpopulation and how this changes through time. This type of information has important implications for determining habitat management targeted at different subpopulations (e.g., residents vs migrants) and for identifying critical habitat types that are being used by all members of a population. Our integrated framework allows us to estimate the degree to which factors that differ across space and time may disproportionately affect a given subpopulation. For example, when assessing the environmental impact of a wind energy development project, we would like to be able to differentiate the potential impact to year-round resident eagles separately from migratory individuals. Similarly, we can generate information that is critical for the field of migration ecology, such as estimates of the spatial distribution of a given population of birds that originated from a specified starting location as they progress through the spring migration. This type of fine-scale, connectivity information has been shown to be biased if based solely on tracking data, without any additional data sources (Rushing et al. 2021). These predictions of migratory pathways consider the estimated spatial proportion of birds in each of the two modeled subpopulations, and the movement dynamics of each subpopulation, given that birds start in the marked location. This information is critical for environmental impact assessments, such as predicting the spatial overlap with proposed land use changes for individuals that may not normally be considered residents of the area of interest. Predictions such as this can help practitioners and managers obtain a more comprehensive measure of the spatio-temporal region that is relevant for wildlife under their jurisdiction, as it provides quantified predictions of where birds that winter in user-defined spatial regions could be over the course of a migratory season. These are just some examples of information that are critical for wildlife conservation and management that can only be obtained by using an integrated model or are improved by jointly using both individual and species distribution data.

Here we focus solely on movement during the spring migration. Extending our modeling approach to the full annual cycle of a migratory species is possible but would require that our model be expanded to allow attractive points of each animal 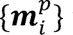 to change over time, to coincide with migratory, wintering, and summering behavior. Extensions to our model such as this are highly feasible given that our modeling framework is very computationally efficient. However, achieving this extension would require additional model development to adequately capture the timing and variation of migration across a species’ range.

Similarly, in our case study we only considered the case where we had two pre-defined subpopulations and pre-assigned each individual bird-year of tracking data to one of the subpopulations. This captured the most obvious variation in migratory patterns of the data we considered here, but for many other scenarios it would be more realistic to consider both a larger number of subpopulations and to model each bird’s assignment to one subpopulation stochastically. Selection of one of several plausible models, each with a different number of subpopulations, could be done by using information criteria such as WAIC or DIC (Hooten and Hobbs 2015), and placing a prior on each bird’s subpopulation assignment would result in a straightforward mixture model extension of our existing model. As the number of subpopulations increases, identifiability of subpopulations could be challenging, and we recommend that researchers contemplating such analyses consider prior distributions that help identify different subpopulations, for example, by ordering subpopulations by overall size, as is commonly done in Bayesian nonparametric statistics. In addition, we modeled each bird year as independent of all other bird years. However, to capture correlation among bird years from the same bird, one could consider a time series prior on the attractive points 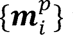, with correlation between successive years.

Independent of potential extensions, the framework we propose leverages existing data and methodologies across disciplines to make unique and robust inferences, allowing us to scale individual movement behaviors to the population level. This approach has the potential to fill critical information needs in the field of migration ecology, but most importantly, provide information critically needed to help conserve and manage species of wildlife that are declining worldwide.

## Acknowledgements

FEB was supported by the U.S. Department of Agriculture National Institute of Food and Agriculture and Hatch Appropriations under Hatch Project #PEN04758 and Accession #1024904. EMH was supported by NSF DMS-2015273. RKM is per contract with the U.S. Fish & Wildlife Service - Division of Migratory Bird Management, National Raptor Program.

## Data Availability Statement

eBird Status and Trends data are publicly available. Golden eagle location data, acquired via satellite telemetry, may be accessed by request to the U.S. Fish and Wildlife Service’s Division of Migratory Bird Management, National Raptor Program. Details on the MCMC algorithm for fitting the model and a validation simulation are presented in a supplemental file. At this time, simulated data and code to fit the model can be requested by emailing the corresponding author.

