## Supplement for "Integrated Movement Models for Individual Tracking and Species Distribution Data"

**Appendix A: Trace Plots and Details of Computing**

As described in the main manuscript, the Integrated Movement Model (IMM) is fit to existing species distribution data using a Markov chain Monte Carlo algorithm. We provide some details of this algorithm, and trace plots for some parameters, in this appendix. There are 26 parameters to be estimated in the IMM, 12 for each subpopulation and two parameters for the population level species distribution model. The MCMC algorithm is written in R, and code to implement the algorithm is found in the code supplement. The MCMC algorithm works by doing a joint Metropolis-Hastings (MH) update for all 12 subpopulation parameters for subpopulation 1, followed by a joint Metropolis-Hastings (MH) update for all 12 subpopulation parameters for subpopulation 2, followed by a joint Metropolis-Hastings (MH) update for the 2 remaining parameters: $N$ from the multinomial model for the species distribution data (eqn. 16) and $w_1$ – the percent of the population in subpopulation 1 (eqn. 15). After these MH-updates are done, the movement centers $\{\boldsymbol{m}_i^p\}$ for each individual animal are updated one at a time using conjugate updates.

**Appendix B: Simulation Example**

Following DiRenzo et al. (2022), we conduct a basic validation simulation example to illustrate identifiability of our model parameters and soundness of the custom code used to fit the model. We first fixed all parameters in the IMM to their posterior mean values. To simulate telemetry data, we simulated the same number of individuals from each population as are present in our golden eagle data, and for each animal we simulated starting at the first recorded data point (each simulated bird begins at the first recorded data point for an observed bird). All other time points were simulated from the IMM.

We simulated species distribution data directly from the IMM, by first finding the posterior mean distribution of each subpopulation at each weekly time point, and then drawing directly from a multinomial distribution (eqn 16) with mean defined by the posterior mean subpopulation distribution. The simulated species distribution data is then the weighted sum of the subpopulation multinomial random variables, with weighting defined by the posterior mean of $w_1$.

These two simulated data streams were then used to fit the IMM, with identical priors as used for the golden eagle analysis. Plots of the posterior distributions of model parameters in this simulation example are shown below. These posteriors show that almost all true parameters used for simulation are reasonably within the posterior distributions, and those parameters which fall into the tails of the posterior are still close to the true values in terms of the absolute numeric difference. Following DiRenzo et al. (2022), we conclude that this simulation example shows that there are no major identifiability issues with our model, and that our code is relatively sound.

Code to replicate this simulation study is found in the code supplement.

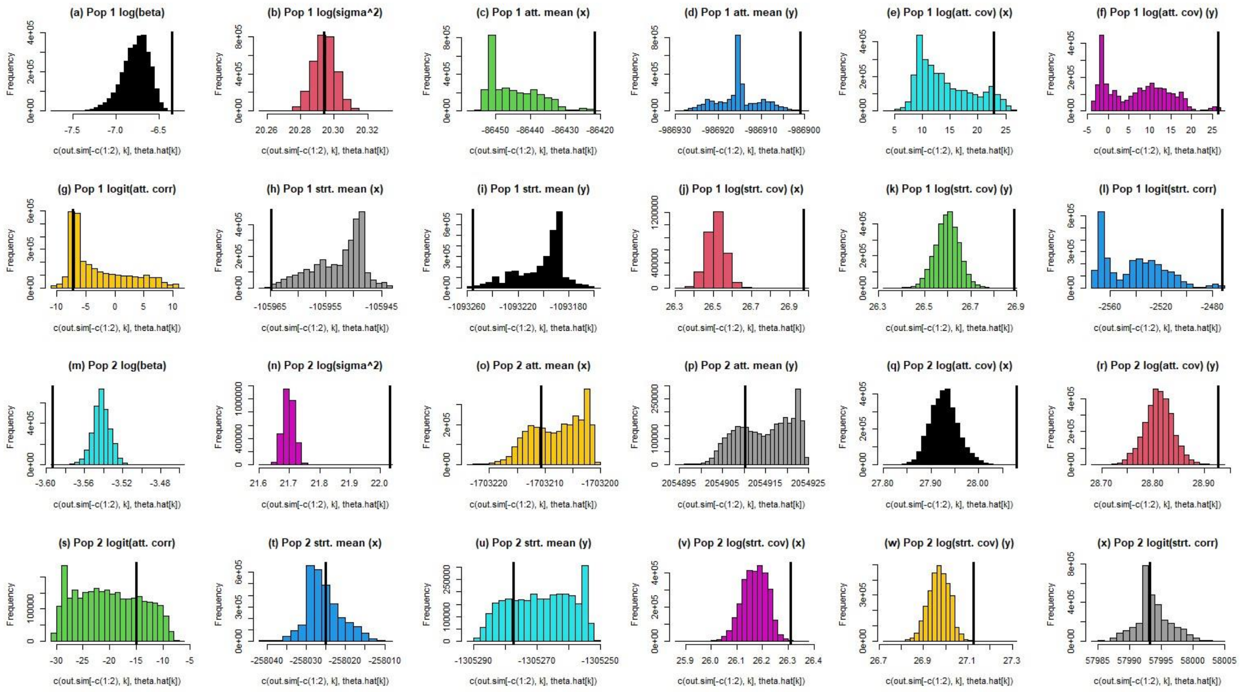


Figure B1: Posterior histograms for all parameters in a simulation study. Vertical black lines indicate the true values used for simulation. The majority of the posterior distributions overlap the true value use to simulate data. Those true values that are outside the bulk of the posterior distributions are qualitatively close to the true values and produce qualitatively very similar behavior.
